# Visual Analysis of Multi-Omics Data

**DOI:** 10.1101/2024.04.23.590648

**Authors:** Austin Swart, Ron Caspi, Suzanne Paley, Peter D. Karp

## Abstract

We present a tool for multi-omics data analysis that enables simultaneous visualization of up to four types of omics data on organism-scale metabolic network diagrams. The tool’s interactive web-based metabolic charts depict the metabolic reactions, pathways, and metabolites of a single organism as described in a metabolic pathway database for that organism; the charts are constructed using automated graphical layout algorithms.

The multi-omics visualization facility paints each individual omics dataset onto a different “visual channel” of the metabolic-network diagram. For example, a transcriptomics dataset might be displayed by coloring the reaction arrows within the metabolic chart, while a companion proteomics dataset is displayed as reaction arrow thicknesses, and a complementary metabolomics dataset is displayed as metabolite node colors. Once the network diagrams are painted with omics data, semantic zooming provides more details within the diagram as the user zooms in. Datasets containing multiple time points can be displayed in an animated fashion. The tool will also graph data values for individual reactions or metabolites designated by the user. The user can interactively adjust the mapping from data value ranges to the displayed colors and thicknesses to provide more informative diagrams.

## 1 Introduction

Scientists are faced with a coming deluge of single- and multi-omics datasets, such as transcriptomics data plus metabolomics data, reaction flux measurements plus transcriptomics data, and transcriptomics data plus proteomics data. Although these data hold great promise, they present sizable analysis challenges; data analysis is a major bottleneck to discovery from multi-omics technologies. Developing computational methods to extract new understanding from complex multi-omics data presents significant challenges in terms of conveying the aspects of a biological system that are changing, and facilitating comparisons of the different omics measurements for the same biological subsystem.

We present a tool for multi-omics data analysis that enables simultaneous visualization of up to four types of omics data on organism-scale metabolic charts. The tool is an expansion of an earlier tool called the Cellular Overview [1, 2]. The Cellular Overview is a web-based interactive metabolic chart that depicts the metabolic reactions, pathways, and metabolites of a single organism as described in a metabolic pathway database for that organism. The tool generates its organism-specific metabolic charts using automated graphical layout algorithms and provides semantic zooming of its diagrams. The tool was quite popular for analysis of single-omics datasets because it directly conveys to the user the changes in activation levels of different metabolic pathways in the context of the full metabolic network.

This article reports significant multi-omics extensions to the Cellular Overview. The tool can now read a multi-omics dataset from a single file or from a set of files. That multi-omics dataset can contain up to four single-omics datasets. Each of those datasets specifies on which of four “visual channels” within the Cellular Overview the dataset will appear. Those channels are the color and thickness of reaction edges within the diagram, and the color and thickness of metabolite nodes within the diagram. For example, a multi-omics dataset could depict transcriptomics data as the color of the metabolic-reaction edges within the diagram, proteomics data as the thickness of reaction edges, one type of metabolomics data as the color of metabolite nodes, and a second type of metabolomics data (possibly measured using a different technology) as the thickness of metabolite nodes. Each of the visual channels can be animated, with manual stepping of the animation if desired. The user can precisely control the mapping from the color assignments and thicknesses within the diagram to the associated data values. The user can also magnify regions of the display and graph sequences of data values.

The multi-omics Cellular Overview is part of the larger Pathway Tools (PTools) software system [3, 4]. PTools is an extensive bioinformatics software system whose capabilities include genome informatics, pathway informatics, omics data analysis, comparative analysis, and metabolic modeling. Pathway informatics features include metabolic reconstruction, pathway search, pathway visualization, and metabolic route search. Thus, the metabolic pathway databases on which the Cellular Overview omics-data analysis tools depend can be created through the metabolic reconstruction component of PTools. PTools is used in a wide spectrum of life-science applications, facilitating studies of sequenced bacteria, plants, and animals, and enabling metabolic engineering. No other software system known to us enables a user to install one software tool to access such a large number of seamlessly integrated capabilities. PTools contains another multi-omics analysis tool called the Omics Dashboard [5, 6] that provides a hierarchical model for analyzing multi-omics datasets; the multi-omics Cellular Overview provides an alternative, metabolism-centric approach to analyzing multi-omics data.

The remainder of this article discusses related work on metabolic network-based omics visualization. It describes in more detail what we mean by a multi-omics dataset, and the file formats in which the tools accept multi-omics data. We then describe the visualizations produced by the multi-omics Cellular Overview and the controls provided with the diagram.

## 2 Related Work on Metabolic Network-Based Omics Visualization

Here we compare the PTools Cellular Overview with related tools for visual pathway-based analysis of omics data. The comparison considers several dimensions of these tools: Do they support analysis of multi-omics data, and how? How are their underlying diagrams produced, e.g., must the metabolic network diagrams be drawn manually (which is quite time consuming), or are they produced automatically? Do the underlying diagrams support important functionality such as semantic zooming and animated displays?

Three approaches have been used to produce drawings of large metabolic networks, independent of their use for analysis of omics data. Approach 1: General graph layout algorithms have been used by Cytoscape [7] and VisANT [8]. Diagrams produced by these algorithms bear little resemblance to textbook pathway diagrams, which means these diagrams are less familiar to biologists and therefore more time consuming to learn. Such diagrams are also less useful because creators of biological pathway drawings have adopted specialized graphical conventions for a reason — they speed the understanding of pathways. Approach 2: Manually drawn pathways have been used by KEGG-related tools and Escher. Although this type of diagram uses biological pathway drawing conventions, the only way to scale this approach to thousands of genomes is to create large “uber pathway diagrams” that combine pathways from many organisms, and thus a single uber diagram can be re-used across many organisms. However, such diagrams necessarily contain many pathways that are not present in any one organism, making the diagrams confusing and larger than necessary. Furthermore, these diagrams must be updated manually when the underlying pathway DB is updated. Approach 3: Pathway-specific layout algorithms have been used by PTools. This approach produces organism-specific diagrams containing just those pathways present in a given organism. This approach scales to thousands of organisms and can be updated automatically so that the diagram always reflects the latest version of each pathway. These algorithms are difficult to develop.

Table 1 compares omics pathway-painting tools along the following six dimensions. (1) Do their diagrams use general layout algorithms, manual uber drawings (MAN), or pathway-specific algorithms (PSA)? (2) Do they paint omics data on whole metabolic network diagrams? Some tools paint omics data onto single-pathway diagrams only. (3) How many omics datatypes can the tool visualize simultaneously? (4) Can the tool depict omics pop-ups that graph omics data for a given gene or metabolite, which provide more precise quantitative information than the colored grid squares that many diagrams produce? (5) Can the tool produce animated displays to depict multiple time points or conditions? (6) Does the whole-network diagram support semantic zooming that provides more information as the user zooms in, such as gene and metabolite names?

**Table 1:** Comparison of omics painting tools.

| Capability | PTools | KEGG Mapper | PathView Web | Paint Omics | KEGG Atlas | iPath | Escher |
| --- | --- | --- | --- | --- | --- | --- | --- |
| (1) Diagram Type | PSA | MAN | MAN | MAN | MAN | MAN | MAN |
| (2) Full Network Omics | YES | YES | no | no | YES | YES | YES |
| (3) Simultaneous Omics | 4 | 2 | 2 | 3 | ? | 2 | 2 |
| (4) Omics Popups | YES | no | no | no | no | no | no |
| (5) Animation | YES | no | no | no | no | YES | no |
| (6) Semantic Zooming | YES | no | no | no | no | no | no |

Table 1 provides information about the following software tools: PTools [4] and KEGG Mapper [9], which are the only tools that paint data onto both full metabolic network diagrams and individual metabolic pathways. PathView Web [10] and PaintOmics 3 [11] paint multi-omics data onto single-pathway diagrams, not onto full metabolic network diagrams; the same is true of BioMiner [12] (not listed in table — defunct, website not active). KEGG Atlas [13] (defunct) and iPath 2.0 [14] paint data onto full metabolic network diagrams. Escher [15] paints data onto any network diagrams manually created by the user, which can be for individual pathways, collections of pathways, or full metabolic networks.

PTools is the most powerful tool, providing organism-specific metabolic-network diagrams that are generated automatically, support semantic zooming, and enable visualization of up to four omics datasets simultaneously with animation and omics popups.

## 3 Multi-Omics File Formats

We consider a multi-omics dataset that can be analyzed using the Cellular Overview to be a collection of from one to four single-omics datasets. The single-omics datasets can be measurements of transcript abundance (e.g., transcriptomics), protein abundance (e.g., proteomics), reaction flux, or metabolite abundance (e.g., metabolomics). For that matter, a single-omics dataset can contain any numeric values that the user wants to map to the reaction edges or metabolite nodes of the metabolic network diagram. Each of these single-omics datasets can contain one or more time points or conditions, which are represented as columns in the corresponding file. It is preferable, but not required, that each single-omics dataset contain the same number of columns (time points or conditions).

A multi-omics dataset can be provided to the Cellular Overview in two ways: as a single file and as a set of files, one per single-omics dataset. This approach provides flexibility both to those users who prefer to separate out each single-omics dataset into a separate file, and to those users who prefer to put all their data into one file. Another design goal for the file format is to capture the settings of parameters for the Cellular Overview so that the user can store these parameters into the file once, rather than having to enter them every time they invoke the Cellular Overview.

### 3.1 Single-File Multi-Omics Format

This format enables up to four single-omics datasets to be combined together into one text file. The file is organized with a header section at the beginning that specifies how many single-omics datasets will follow. The header also defines Cellular Overview parameters for each dataset including which data columns from the single-omics dataset should be used, the type of data in that dataset (e.g., “gene” versus “compound”), an identifying text label for that dataset, and a unique identifier for that dataset (e.g., “Table1”).

Next, there is a file section for each single-omics dataset. Each single-omics dataset is encoded as a table consisting of a set of rows and columns. Each row describes a single gene, protein, metabolite, etc., depending on the type of data in that single-omics dataset. For example, for transcriptomics data, the first column in each row is the name or identifier of a gene. Subsequent columns, which are tab-delimited, list numeric data values for the transcriptomics measurements at different time points or conditions.

An example file in single-file format is provided as Supplemental File 1. The file format is documented online in Section 9.3.7 of [16].

### 3.2 Multi-File Multi-Omics Format

The multi-file format separates the sections within the single-file format into multiple files. The header section from the single-file format becomes a separate file called the master file. The master file is where parameter values are assigned for each omics dataset, and the master file lists the file names for each single-omics dataset. Each data table section from the single-file format is stored in a separate file.

## 4 Multi-Omics Cellular Overview Visualizations

This section demonstrates the use of the Multi-Omics Cellular Overview on multi-omics datasets for *Acinetobacter baylyi* and *Synechocystis* PCC 6803.^1^

### 4.1 Single Time Point Dataset for *Acinetobacter Baylyi*

Figure 1 depicts a multi-omics dataset using the Cellular Overview. The diagram is organized to show metabolic pathways and reactions inside the cell membrane, with individual reactions on the right side, and pathways on the left. Pathways flow downwards. The diagram is divided so that biosynthetic pathways are drawn on the left portion of the pathway area and catabolic pathways are on the right side. Further subdivisions group pathways into categories such as cofactor biosynthesis and carbohydrate degradation. Edges (lines) in the diagram represent reactions, and nodes represent metabolites. Node shapes communicate the type of metabolite, e.g., triangles are amino acids and squares are carbohydrates; shaded nodes are phosphorylated compounds. As the user zooms in and out of the diagram with a mouse wheel or trackpad, the names of pathways, metabolites, genes, and enzymes appear. The menu to the right (not shown) enables the user to search the diagram by gene name, metabolite name, enzyme name, etc.

**Figure 1.**
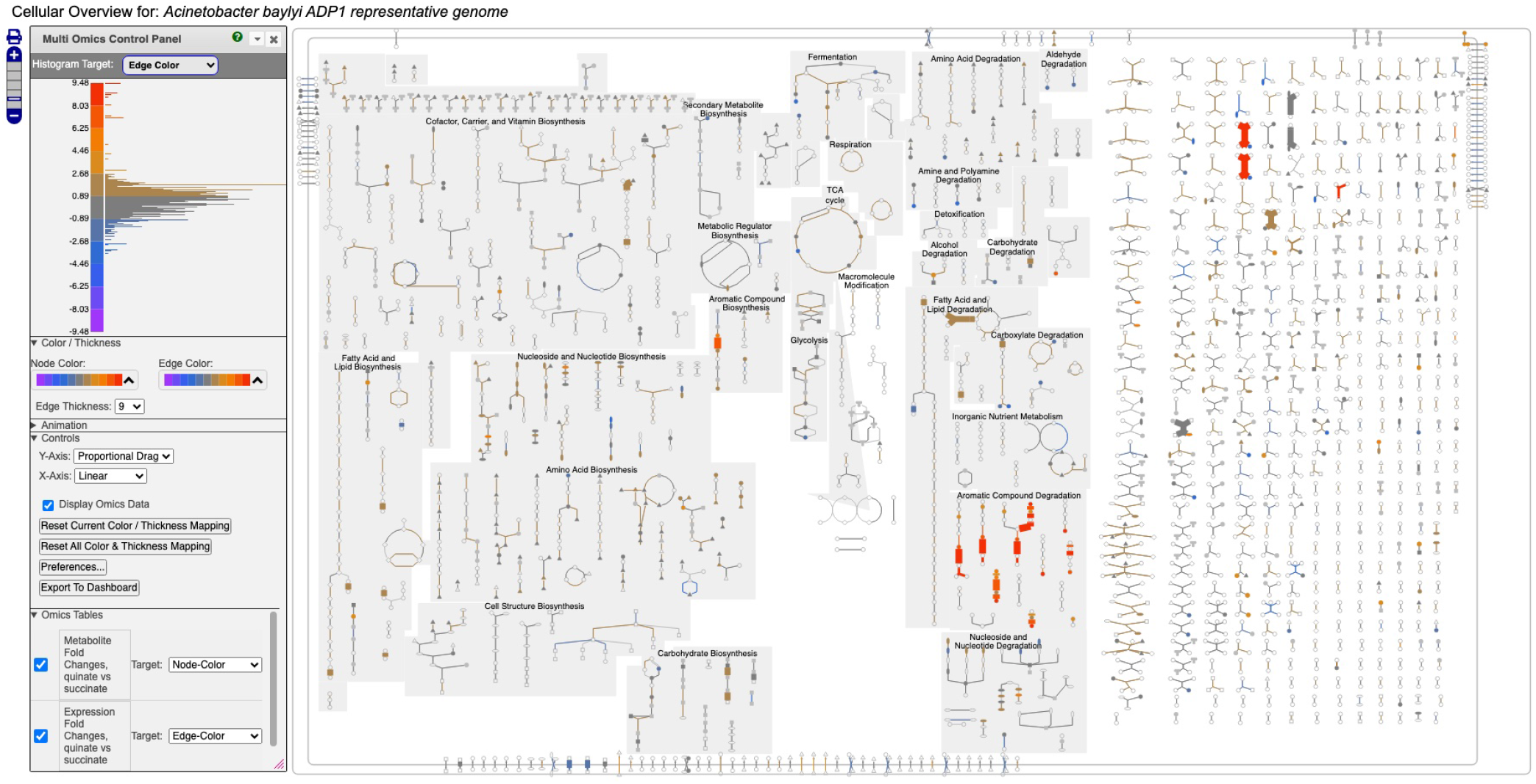
Full Cellular Overview for *A. baylyi* painted with multi-omics data. On the left is the control panel for the Cellular Overview. The histogram at the top of the control panel shows the assignment of data value ranges to edge colors; red and orange colors represent data values from 4.48–9.48.

Figure 1 depicts a multi-omics dataset for *Acinetobacter baylyi* ADP1 in which growth of *A. baylyi* was compared under succinate versus quinate as the carbon source [17]. The component single-omics datasets were (1) a transcriptomics dataset, (2) a gene essentiality dataset, and (3) a metabolomics dataset. All three datasets had a measurement at one time point only. The input data files specified that dataset (1) should be targeted to edge colors, dataset (2) should be targeted to edge thicknesses, and dataset (3) should be targeted to node colors.

Follow these steps to recreate Figure 1:

1. Visit BioCyc.org in a web browser.
2. Click the Change Current Database button and type Acinetobacter baylyi ADP1.
3. Run this command from the top menu: Tools *→* Metabolism *→* Cellular Overview.
4. Run this command from the right-sidebar operations menu: Upload Multi-Omics Data from File.
5. Click the Choose File button in the dialog.
6. Provide Supplemental File 1 as the input file.
7. Click the Submit button in the dialog.

Figure 1 shows the resulting Cellular Overview diagram painted with the preceding multi-omics dataset. Although changes are evident in many areas of metabolism, they are particularly strong in the area Aromatic Compound Degradation, which contains many red nodes and edges and lies to the bottom-right of center. Zooming in on that region initially presents the diagram in Figure 2, which shows that metabolite changes (node colors), gene expression changes (edge colors), and gene expression changes (edge thicknesses) are well coordinated (they are all high). As we further zoom Figure 3, metabolite shapes (e.g., circular nodes) are replaced by metabolite names; the names are colored with omics data. The new carbon source, quinate, appears in a pathway near the middle of the figure, quinate degradation I. The end product of that pathway is protocatechuate, which is the input to the pathway at the top center of Figure 3, namely aromatic compounds degradation via beta-ketoadipate. Thus, the Cellular Overview has clearly shown the pathways that the cell activates to catabolize quinate.

**Figure 2.**
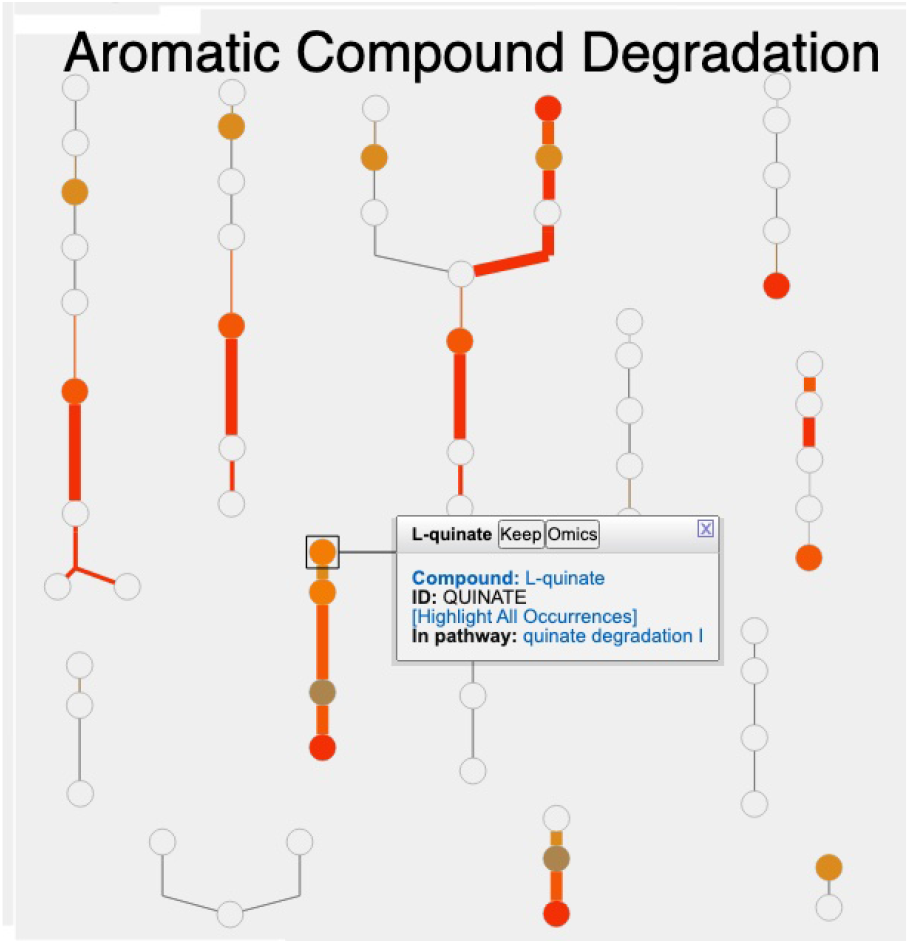
Visualization of *A. baylyi* multi-omics data on the Aromatic Compound Degradation region of the Cellular Overview; the remaining regions of the Cellular Overview have been cropped away.

**Figure 3.**
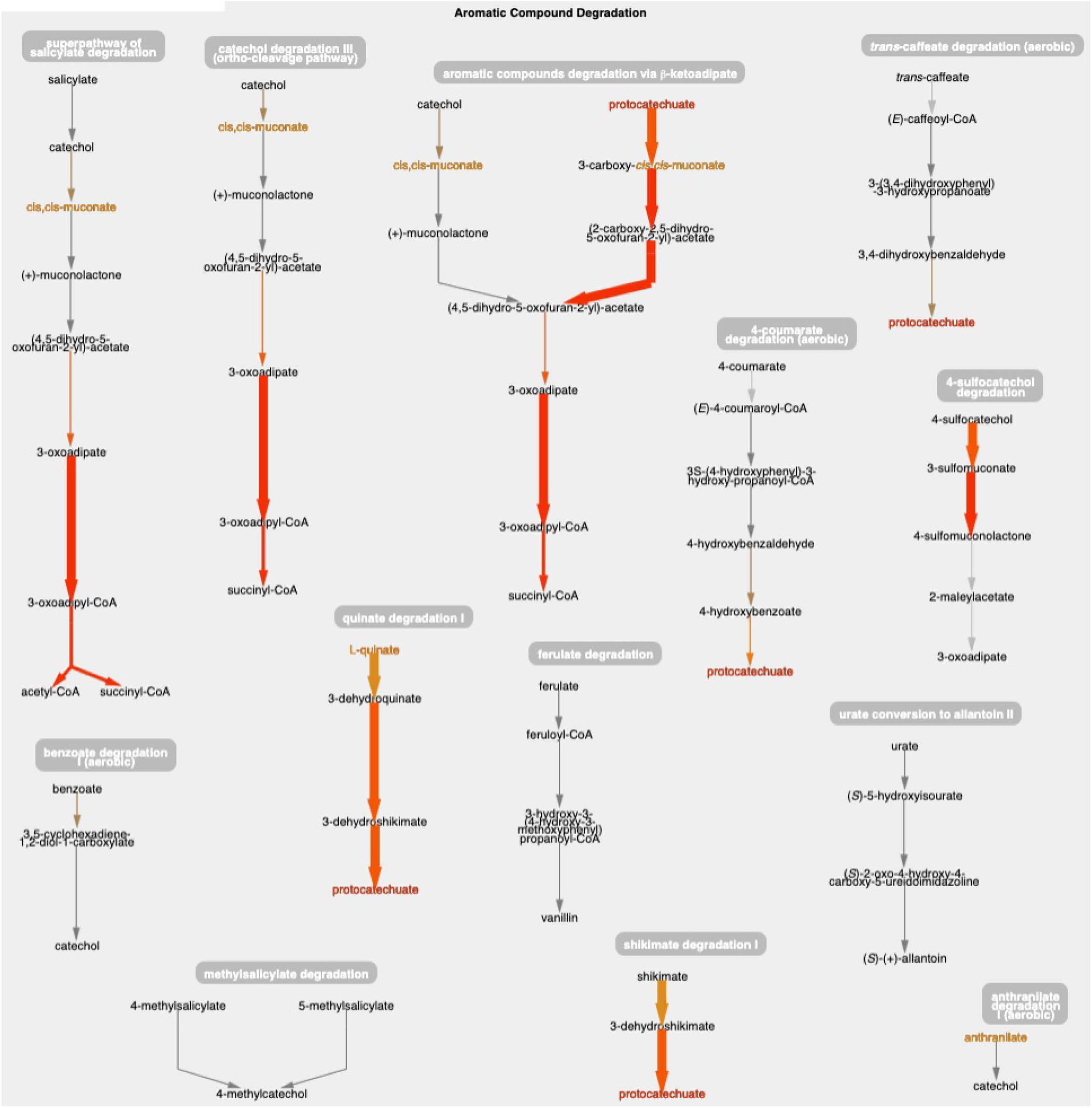
The Aromatic Compound Degradation region of the *A. baylyi* Cellular Overview zoomed to a higher magnification than Figure 2, causing metabolite nodes to become visible.

The only two amino acids whose abundances are increased during growth on quinate are the aromatic amino acids tyrosine and phenylalanine, which can be found in the Amino Acid Biosynthesis region of the Cellular Overview. [17] noted this fact, and postulated that products of quinate degradation in the periplasm may be leaking into the cytoplasm where they become inputs to the biosynthetic pathways for these two amino acids.

### 4.2 Multiple Time Point Dataset for *Synechocystis* PCC 6803

Figure 4 depicts a multi-omics dataset for the cyanobacterium *Synechocystis* sp. PCC 6803 from a study of the effects of the transition from light to dark on this organism [18]. Five time points were taken at 1 hr, 3 hr, 11.75 hr (right before the transition), 13 hr, and 15 hr. The authors reported transcriptomics and proteomics data at all time points. They also provided limited metabolomics data consisting of a list of the most changed metabolites, with a corresponding score for each, but no time points.

**Figure 4.**
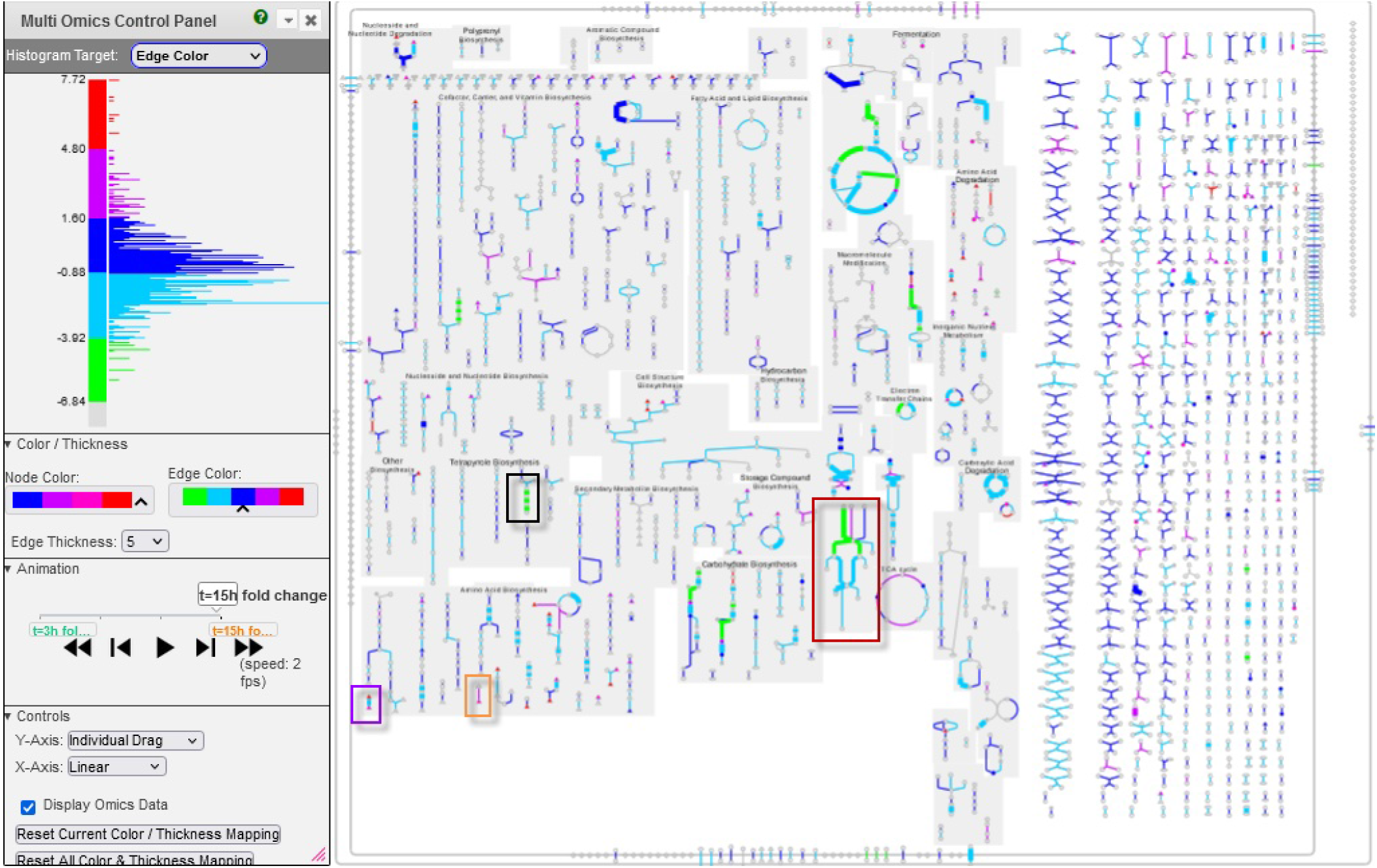
Multi-time point dataset for *Synechocystis* PCC 6803 shown on the Cellular Overview. The omics viewer control panel is on the left side of the diagram. In this diagram, edge color is assigned to transcriptomics data, edge thickness is assigned to proteomics data, and node color is assigned to metabolomics data. The histogram shown in the image describes the color distribution for depicting transcriptomics data. The red, purple, orange, and black rectangles point to data described further in Figures 5, 6, 7, and 8, respectively.

We generated the input file (Supplemental File 2) used to create Figures 4–8 by computing fold changes relative to the 1 hr timepoint, so that transcriptomics and proteomics datasets each consist of 4 datapoints. Only one time point is available for the metabolomics data, consisting of the reported scores. The input file specifies that the transcriptomics dataset should be targeted to edge colors, the proteomics dataset should be targeted to edge thicknesses, and the metabolomics dataset should be targeted to node colors.

Follow these steps to recreate Figure 4:

1. Visit BioCyc.org in a web browser
2. Click the Change Current Database button and type “Synechocystis sp. PCC 6803 substr. Kazusa”
3. Run this command from the top menu: Tools *→* Metabolism *→* Cellular Overview
4. Run this command from the right-sidebar operations menu: Upload Multi-Omics Data from File
5. Click the Choose File button in the dialog
6. Provide Supplemental File 2 as the input file
7. Click the Submit button in the dialog

Depending on your selection of color palettes for the different datasets and the ranges you specify for the histograms, your image may differ in appearance from Figures 4–8.

Click on the play button to start the animation. The diagram will move through the time points, showing significant changes in many of the pathways and reactions.

We now present several examples of where using this tool can help users to observe interesting phenomena found in these omics data.

#### Example 1

Using the animation feature, the tool makes it easy to see multiple changes that occur simultaneously. For example, Figure 5 shows the “superpathway of electron transport in the thylakoid membrane” at the latest time point (towards the end of the dark period). One can see how photosystem II (green line) has low transcription (green color) and high protein concentration (thick line) at this time point. At the same time, transcription of the succinate dehydrogenase complex (purple line) and Ndh2 (NADH-quinone oxidoreductase) is significantly higher (no proteomic data is available for these two complexes), indicating that, during the dark, the main sources for reducing power are succinate and NADH.

**Figure 5.**
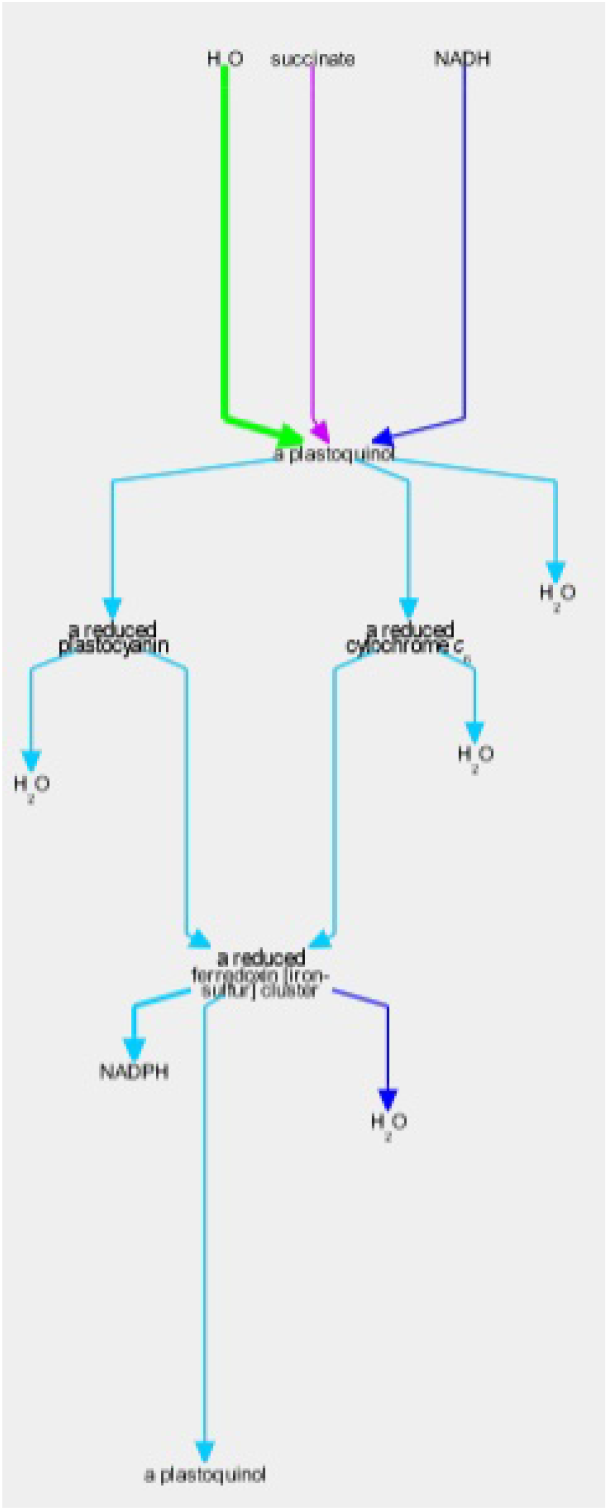
The *Synechocystis* pathway “superpathway of electron transport in the thylakoid membrane” extracted from Figure 4, at the latest time point. The pathway combines several electron transport pathways that occur in the thylakoid membrane, including components of linear photo-synthetic electron transport, respiratory electron transport, cyclic electron transport, and photo-protection by flavodiiron proteins.

#### Example 2

Glutamate and glutamine are among the most dynamic metabolites in this study [18]. They commented that GlnA (glutamine synthetase type I) shows strong downregulation in the dark, while GlnN (glutamine synthetase type III) protein levels showed no significant change. By hovering the mouse on the glutamine synthetase reaction in the “L-glutamine biosynthesis” pathway and selecting “Omics,” a popup diagram appears that clearly illustrates this observation (Figure 6).

**Figure 6.**
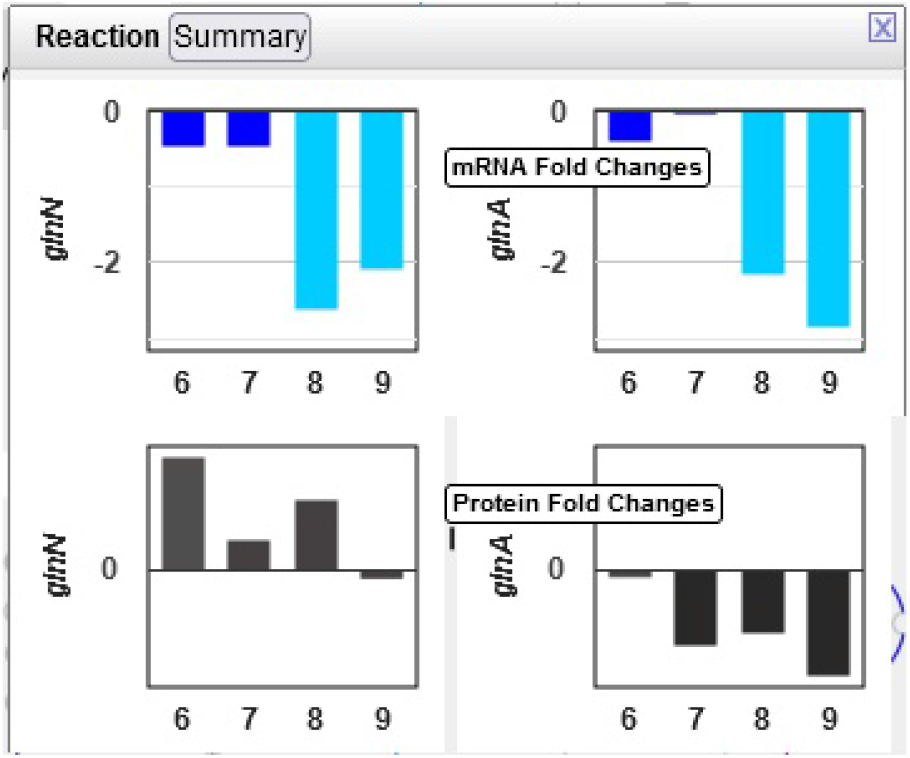
Omics pop-ups graphing *Synechocystis* transcriptomics and proteomics data for the *glnA* and *glnN* genes and their products, which catalyze the ATP-dependent conversion of L-glutamate to L-glutamine. The transcription of both genes is strongly down-regulated in the dark, but proteomics data indicate a much stronger decline in the amount of GlnA than of GlnN.

#### Example 3

The only amino acid whose synthesis was enhanced in the dark was L-alanine. The only L-alanine biosynthetic route demonstrated so far in *Synechocystis* is desulfuration of L-cysteine, and three enzymes are known to catalyze this reaction. Opening the relevant popup suggests that the responsible enzyme under these conditions is the cysteine desulfurase encoded by the sll0704 gene (shown in Figure 7 as SGL RS02380, the accession ID assigned by RefSeq).

**Figure 7.**
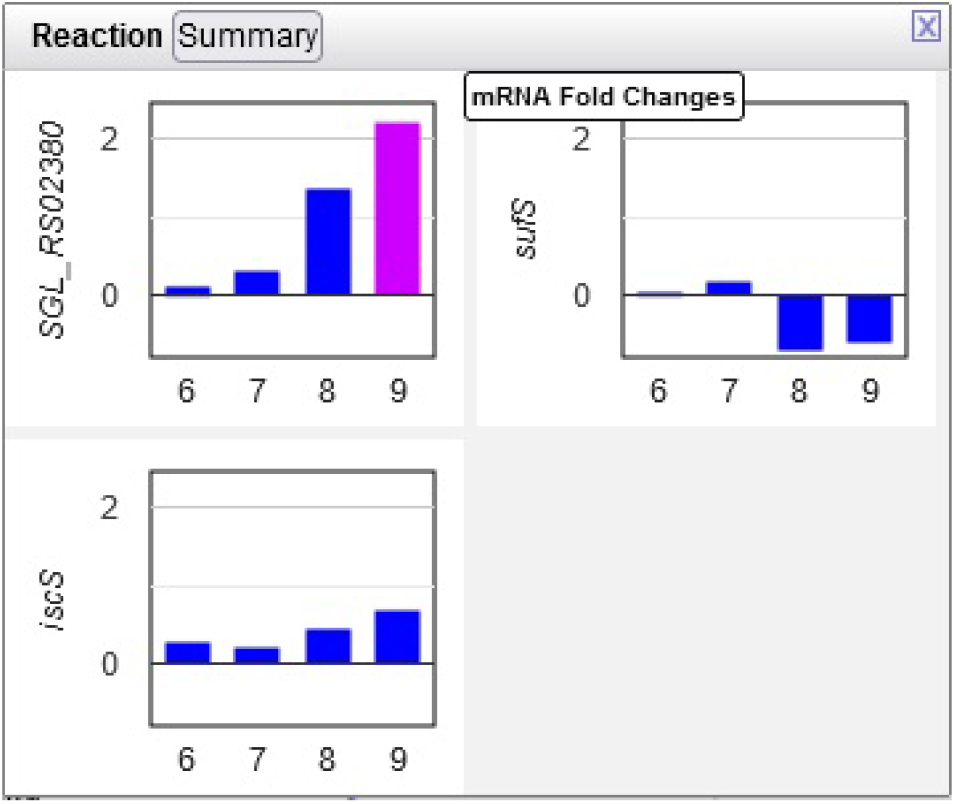
Transcriptomics data for three *Synechocystis* cysteine desulfurases. Transcription of the sll0704 gene (shown as SGL RS02380) is strongly up-regulated in the dark, whereas the transcription of the *iscS* and *sufS* genes does not change significantly.

#### Example 4

The authors noted that chlorophyll content increased in the first three hours of the light period. Looking at the multiomics data for ChlP (Figure 8), the enzyme that converts chlorophyllide a to chlorophyll a and part of the “chlorophyll a biosynthesis II” pathway, one can see a strong delay in the degradation of the enzyme, as the levels of the enzyme continue to grow in the dark despite a very strong decline in transcription.

**Figure 8.**
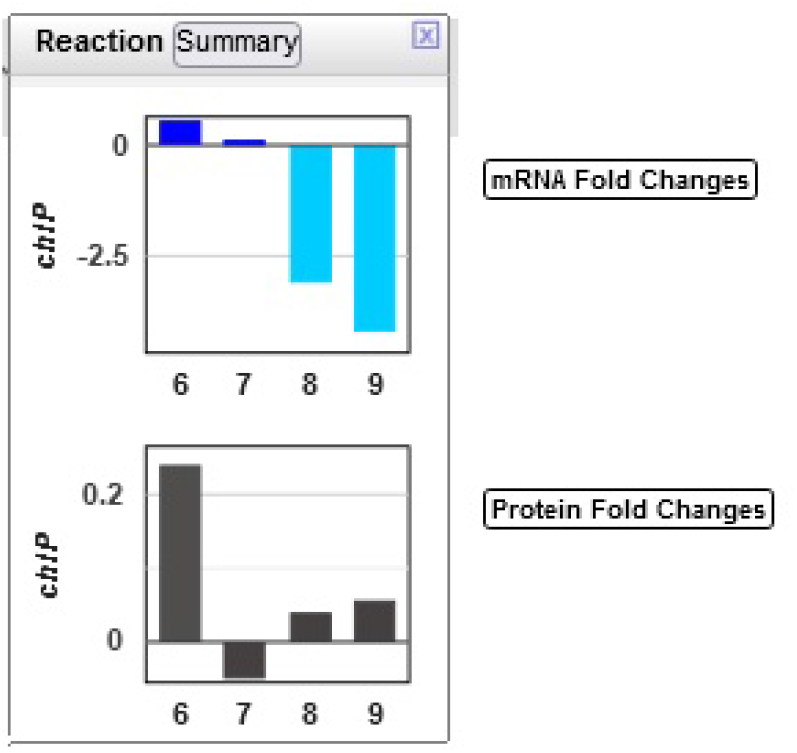
Transcriptomics and proteomics data for the *Synechocystis chlP* gene and its product, geranylgeranyl diphosphate reductase, which catalyzes a reaction during chlorophyll a biosynthesis.

### 4.3 Omics Viewer Controls

Several omics-viewer controls enable the user to tune the diagram to meet their needs more effectively. The controls are shown in the left side of Figure 4.

With the top control, the user can adjust the mapping of colors and thicknesses to data values. Currently the histogram at the top left shows the number of Cellular Overview data points that are mapped to each of the y-axis values. For example, the most data points are in the dark blue and light blue regions that correspond to data values near 0. The histogram is labeled “Histogram Target: Edge Color” to indicate that the diagram is currently depicting edge colors, but the user can also select node colors, node thicknesses, and edge thicknesses.

Currently the control shows that for edge colors there is a linear mapping between data values from -7.72 to 7.72 and the five colors (green to red) shown. If the user wanted the red and green colors at the extremes to encompass a larger set of data values, they could use the mouse to click on the axis label “4.80” and drag that label downwards to expand the length of the red color scale with all of the other regions of the scale decreasing in size. Similarly a user could grab the label “-4.80” and drag it upwards.

With the next control pane, labeled “Color/Thickness,” the user can choose among several provided color scales, each containing a different set of colors. The user can also change the maximum edge thickness the tool uses.

The next pane (“Controls”) enables the user to make other adjustments such as selecting how the y-axis dragging works and resetting the color mapping to the defaults.

The next pane (“Omics Tables,” visible in Figure 1) shows the mapping from each provided single-omics dataset to its target within the Cellular Overview. For example, it shows that expression fold changes are mapped to edge colors. The user can alter the mapping as desired.

If the user mouses over a node or edge within the diagram, a tooltip appears that both identifies that metabolite or reaction and enables the user to display omics pop-ups for that entity and then graph its omics data.

## 5 Conclusions

We have illustrated the use of the extended Cellular Overview multi-omics visualization capabilities using multi-omics datasets for *A. baylyi* and *Synechocystis*. For *A. baylyi,* the tool identified likely pathways that degrade quinate, and confirmed an observation from the original publication of this dataset that two aromatic amino acids increased in abundance. For *Synechocystis,* the tool identified a number of changes in the metabolic state of the organism, including which cysteine desulfurase isozyme is likely to be responsible for the increased level of L-arginine.

The multi-omics Cellular Overview supports coordinated analysis of multi-omics datasets on automatically generated organism-specific metabolic network diagrams. Up to four single-omics datasets can be analyzed simultaneously including transcriptomics, proteomics, and metabolomics data. The datasets are provided via either a single input file or through separate files for each single-omics dataset. The user specifies for each dataset whether it should be displayed as node colors, node thicknesses, edge colors, or edge thicknesses. The user can alter the mapping of data values to the color and thickness scales via interactive sliders. Datasets containing multiple time points can be displayed as animations with the ability to single-step the animations. The tool can also graph the data values for individual nodes and edges.

## 6 Materials and Methods

The software that generates Cellular Overview layouts is written in Common Lisp. The software that displays Cellular Overview diagrams and multi-omics data is written in JavaScript, and has been tested within the Chrome, Firefox, and Safari browsers. Although the functionality described herein can be verified through the interactive BioCyc website, all software described in this article is freely available to research institutions for research purposes as part of the Pathway Tools software, including source code, by request to.

For the example datasets used in this paper, we downloaded the publicly available supplementary data files for [17] and [18]. Where log2 fold change columns were not already present, we computed them (relative to the first timepoint for the *Synechocystis* data, and based on the provided fold change column for the *Acinetobacter* data). The gene essentiality data for the *Acinetobacter* dataset was provided as normalized optical-density growth yields. We subtracted this value from 1 for each gene to generate a measure of essentiality such that higher values represent greater essentiality. The individual component datasets were extracted and combined into a single tab-delimited file for each example, each with an added header section. The resulting files are available as Supplementary Files 1 (the *Acinetobacter* example) and 2 (the *Synechocystis* example).

## Author Contributions

PDK directed the research and authored most of the article. SP and RC constructed the examples and wrote the example sections. AS wrote the software.

## Supplemental Data

Supplemental File 1: Single-file mult-omics dataset for *A. baylyi*.

Supplemental File 2: Single-file mult-omics dataset for *Synechocystis* PCC 6803.

## Footnotes

1 The BioCyc site requires users to create a free account, which provides a one-month free trial, after a number of page views.

